# A simple assay for quantification of plant-associative bacterial nitrogen fixation

**DOI:** 10.1101/2021.03.31.437999

**Authors:** Timothy L Haskett, Philip S Poole

## Abstract

Accurate quantification of plant-associative bacterial nitrogen (N) fixation is crucial for selection and development of elite diazotrophic inoculants that could be used to supply cereal crops with nitrogen in a sustainable manner. Because a low oxygen environment that may not be conducive to plant growth is essential for optimal stability and function of the N-fixing catalyst nitrogenase, quantification of N fixation is routinely carried out on “free-living” bacteria grown in the absence of a host plant. Such experiments may not divulge the true extent of N fixation occurring in the rhizosphere where the availability and forms of nutrients such as carbon and N, which are key regulators of N fixation, may vary widely. Here, we present a modified *in planta* acetylene reduction assay, utilising the model cereal barley as a host, to quantify associative N fixation by diazotrophic bacteria. The assay is rapid, highly reproducible, applicable to a broad range of diazotrophs, and can be performed with simple equipment commonly found in most laboratories that investigate plant-microbe interactions.

**Importance:** Exploiting “nitrogen-fixing” bacteria that reduce atmospheric dinitrogen into ammonia as inoculants of cereal crops has great potential to alleviate current inputs of environmentally deleterious fertiliser nitrogen and drive more sustainable crop production. Accurately quantifying plant-associative bacterial nitrogen fixation is central to the development of such inoculant bacteria, but most assays fail to adequately reproduce the conditions of plant root systems. In this work, we have validated and optimised a simple *in planta* assay to accurately quantify N fixation in bacteria occupying the root and surrounding soil of the model cereal barley. This assay represents a benchmark for quantification of plant-associative bacterial N fixation.

## Introduction

Exploiting diazotrophic bacteria that reduce atmospheric dinitrogen (N_2_) into ammonia (NH_3_^+^) as inoculants of cereal crops has great potential to alleviate current inputs of environmentally deleterious fertiliser nitrogen (N) in agricultural systems to establish more sustainable crop production (1). While a plethora of diazotrophic strains have been isolated that colonise the root compartments (rhizosphere, rhizoplane, and endosphere) of cereals (2), it remains unclear which strains are best suited to agriculture. Elite inoculants should ideally a) competitively colonise and persist in plant root compartments to exert their beneficial effects, b) exhibit some degree of interactive specificity with the target host to prevent promiscuous growth promotion of non-target species, and c) fix and release large quantities of N for assimilation by the plant (3). While no natural bacteria have been categorically demonstrated to satisfy these three criteria, targeted selection and genetic engineering programs are currently underway to assist in the development of elite inoculant strains (2, 4-8).

Quantification of bacterial N fixation is central to the selection and development of elite inoculant strains and is typically carried out using either an acetylene reduction assay (ARA) or ^15^N incorporation assay. ARAs rely on the use of GC-MS to monitor the alternative ability of the N-fixing catalyst nitrogenase to reduce acetylene (C_2_H_2_) to ethylene (C_2_H_4_) (9), whereas ^15^N incorporation assays can be used to track assimilation of fixed ^15^N by bacteria or even by plants following bacterial release of NH_3_^+^ (10-12). Because a low oxygen (< 10%) environment that may not be conducive to plant growth is essential for optimal nitrogenase stability and function (13), both assays are best suited to analysis of bacterial cultures grown in the absence of a host plant. However, this condition may not reflect the true extent of N fixation occurring in the rhizosphere where the availability and forms of nutrients such as carbon (C) and N, which are key regulators of N fixation (5, 14-16), may vary widely. A benchmark assay is needed to assess associative bacterial N fixation in a manner that more accurately reflects conditions in the rhizosphere.

Here, we present a simple *in planta* ARA to quantify N fixation in diazotrophic bacteria occupying the root system of the model cereal barley (Fig 1). We demonstrate that the assay is highly reproducible, rapid, applicable to genetically diverse diazotrophs, and requires minimal equipment commonly found in laboratories investigating plant-microbe interactions.

**Figure 1.**
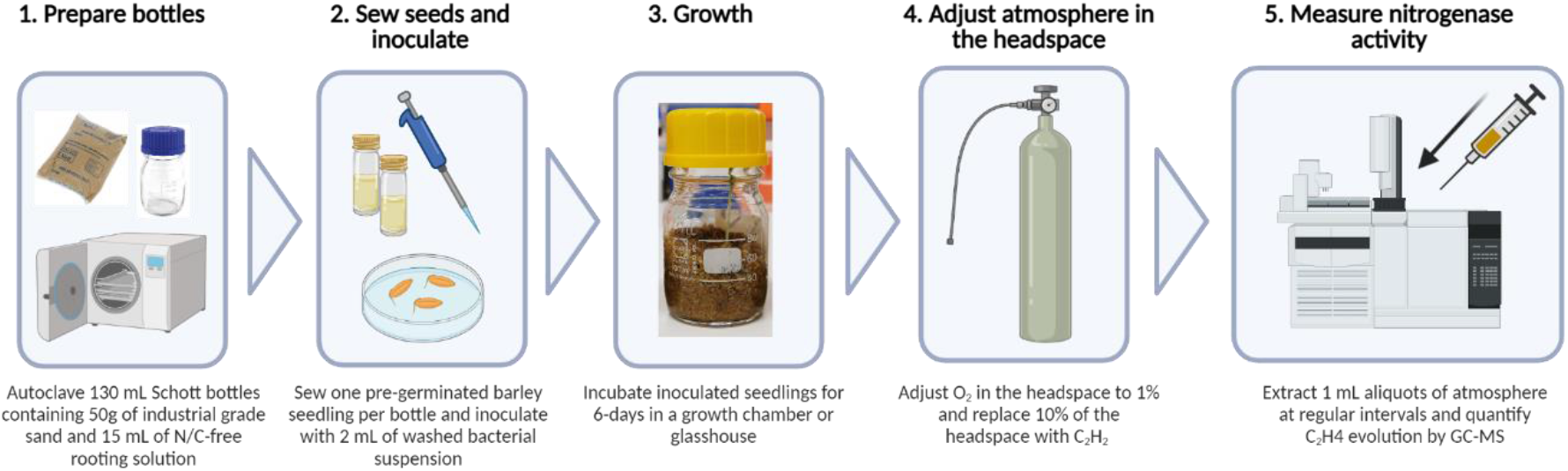
Schematic of workflow for *in planta* acetylene reduction assay. This figure was created with BioRender®.

## Results

### Validation of *in planta* ARA

To both validate and optimise our *in planta* ARA described (Fig 1), we utilised the cereal endophyte and *Sesbania rostrata* nodulating symbiont *Azorhizobium caulinodans* ORS571 (*Ac*) as a model strain. The assay was initially set up by sewing individual pre-germinated, surface-sterilised Barley seeds into 130 mL Schott bottles containing industrial grade sand and N/C-free rooting solution, inoculating the plants with 2 mL of an OD_600nm_ 0.1 suspension of *Ac* (approximately 5×10^7^ cells), then growing the plants in a growth-chamber for 6-days. At this point, the atmosphere in the headspace was adjusted to 1% O_2_ and Schott bottles were sealed with a rubber septum. Ten percent of the headspace was next replaced with C_2_H_2_ and plants were returned to the growth chamber. The reduction of C_2_H_2_ to C_4_H_4_ by nitrogenase was measured over 72-h using GC-MS. In all five biological replicates, significant C_2_H_4_ production (69.97 ± s.e.m 17.05 nmoles C_2_H_4_ h^-1^ plant^-1^) was detected after 24-h and the subsequent rates of C_2_H_4_ production remained stable up to 72-h with a mean rate of nitrogenase activity of 55.57 ± s.e.m. 11.23 nmol C_2_H_4_ h^-1^ plant^-1^ between 24 and 48-h, and 62.77 ± s.e.m. 14.11 nmol C_2_H_4_ h^-1^ plant^-1^ between 48 and 72-h (Fig 2A). Using the above conditions, we also confirmed that neither the plant nor bacteria exhibited nitrogenase activity in the absence of the symbiotic partner between 0 and 72-h (Fig 2B). Thus, bacterial N fixation monitored in the assay was entirely dependent on nutrients provided by the host plant.

**Figure 2.**
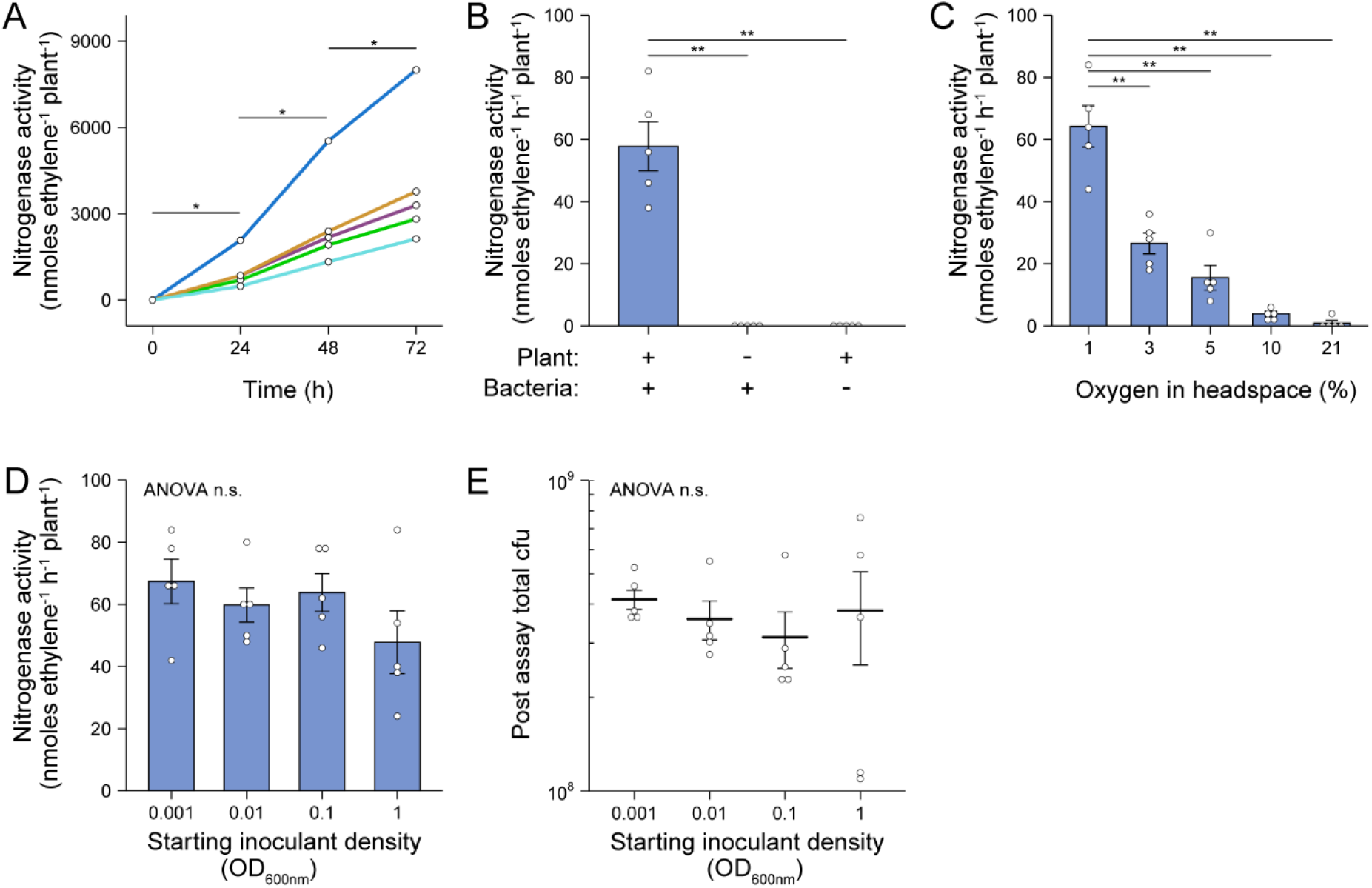
Validation and optimisation of *in planta* acetylene reduction assay. *A. caulinodans* ORS571 (*Ac*) was used as the model strain for these experiments. The assays were set up by sewing individual pre-germinated, surface-sterilised barley seeds into 130 mL Schott bottles containing industrial grade sand and N/C-free rooting solution, inoculating the plants with 2 mL of an OD_600nm_ 0.1 suspension of *Ac*, then growing the plants in a growth-chamber for 6-days. At this point, the atmosphere in the headspace was adjusted to 1% O_2_, the bottles were sealed with a rubber septum and 10% of the headspace was replaced with C_2_H_2_. Bottles were returned to the growth chamber and the reduction of acetylene (C_2_H_2_) to ethylene (C_4_H_4_) by nitrogenase was measured at 24-intervals over 72-h using GC-MS. Mean ± s.e.m (error bars) and individual values for five biological replicates are plotted. Nitrogenase activity was calculated between 24-h and 48-h. ANOVA and pairwise two-tailed students t-tests with Bonferroni adjusted P-values were used to compare means where relevant. P > 0.05 not significant (n.s.), P < 0.05 *, P < 0.01 **, P < 0.001 ***. **(A)** C_2_H_4_ production in each of five biological replicates was monitored over 72-h. **(B)** Rates of nitrogenase activity where measured when the plant or bacteria was omitted from the system. **(C)** Rates of nitrogenase activity were measured when the starting O_2_ concentration in the headspace was adjusted between 1% and 21% (i.e. air) by flushing with N_2_ gas. **(D)** Rates of nitrogenase activity were measured when the starting inoculant density was adjusted between OD_600nm_ 0.001 to 1. **(E)** Total colony forming units (cfu) present in the assay systems of experiment (D) as determined by viable counts after 72-h.

### Optimisation of *in planta* ARA

We further optimised our *in planta* ARA first by titrating the starting O_2_ concentration in the headspace of Schott bottles after 6-dpi. We found that an optimum rate of nitrogenase activity, similar to that of the experiments in Fig 2A-B, was observed where O_2_ in the headspace was adjusted to 1% of the atmosphere (Fig 2C). However, low nitrogenase activity was also observed where the O_2_ concentration was adjusted to 10% (3.98 ± s.e.m. 0.83 nmol C_2_H_4_ h^-1^ plant^-1^) and even 21% (0.89 ± s.e.m. 1.98 nmol C_2_H_4_ h^-1^ plant^-1^), although the latter rate was the product of a single biological replicate out of five (Fig 2C).

We next tested the effect of titrating the starting inoculant density between OD_600nm_ 0.001 and 1.0 and found that nitrogenase activity did not differ between these treatments (Fig 2D). Moreover, when bacteria were recovered at the close of the assay by rigorous flushing with PBS, the total number of colony-forming units (cfu) in each Schott bottle did not differ significantly, with each harbouring between 10^8^ and 10^9^ cfu (Fig 2E). This suggested that after 6-dpi, *Ac* naturally reaches the carrying capacity of the assay system.

### Test of *in planta* ARA on genetically diverse diazotrophic bacteria

To explore whether our *in planta* ARA could be used to quantify associative N fixation by diazotrophs other than *Ac*, we selected the following seven additional alpha-, beta- and gamma-proteobacterial strains for testing; *Azospirillum brasilense* FP2 (*Ab*), *Azoarcus olearius* DQS-4 (*Ao*), *Burkholderia vietnamensis* WPB (*Bv*), *Herbaspirillum seropedicae* SmR1 (*Hs*), *Klebsiella oxytoca* M5a1 (*Ko*), *Pseudomonas stuzeri* A1501 (*Ps*) and *Rhodobacter sphaeoroides* WS8 (*Rs*). Plants were inoculated with 2 mL of an OD_600nm_ 0.01 suspension and after 6-dpi, the atmosphere in the headspace was adjusted to 1% O_2_ and 10% C_2_H_2_ to begin the assay. GC-MS measurements for C_2_H_4_ production were made at 12-h intervals over 72-h. For most of the strains, nitrogenase activity was detectable by 24-h, but a stable, optimal rate of nitrogenase activity required at least 36-h of preparation (Fig 3A). Mean rates of nitrogenase activity were measured for all strains between 48 – 62-h (Fig 3B), with the highest rates between 58 and 65 nmol C_2_H_4_ h^-1^ plant^-1^ for *Ao, Ps, Ac*, and *Ab*. The remaining strains fixed between 7 and 30 nmol C_2_H_4_ h^-1^ plant^-1^, whereas no nitrogenase activity was detected in the uninoculated controls. The range in nitrogenase activity displayed by the eight diazotrophic bacteria tested highlights that some bacteria are better adapted to associative N fixation with barley then are others.

**Figure 3.**
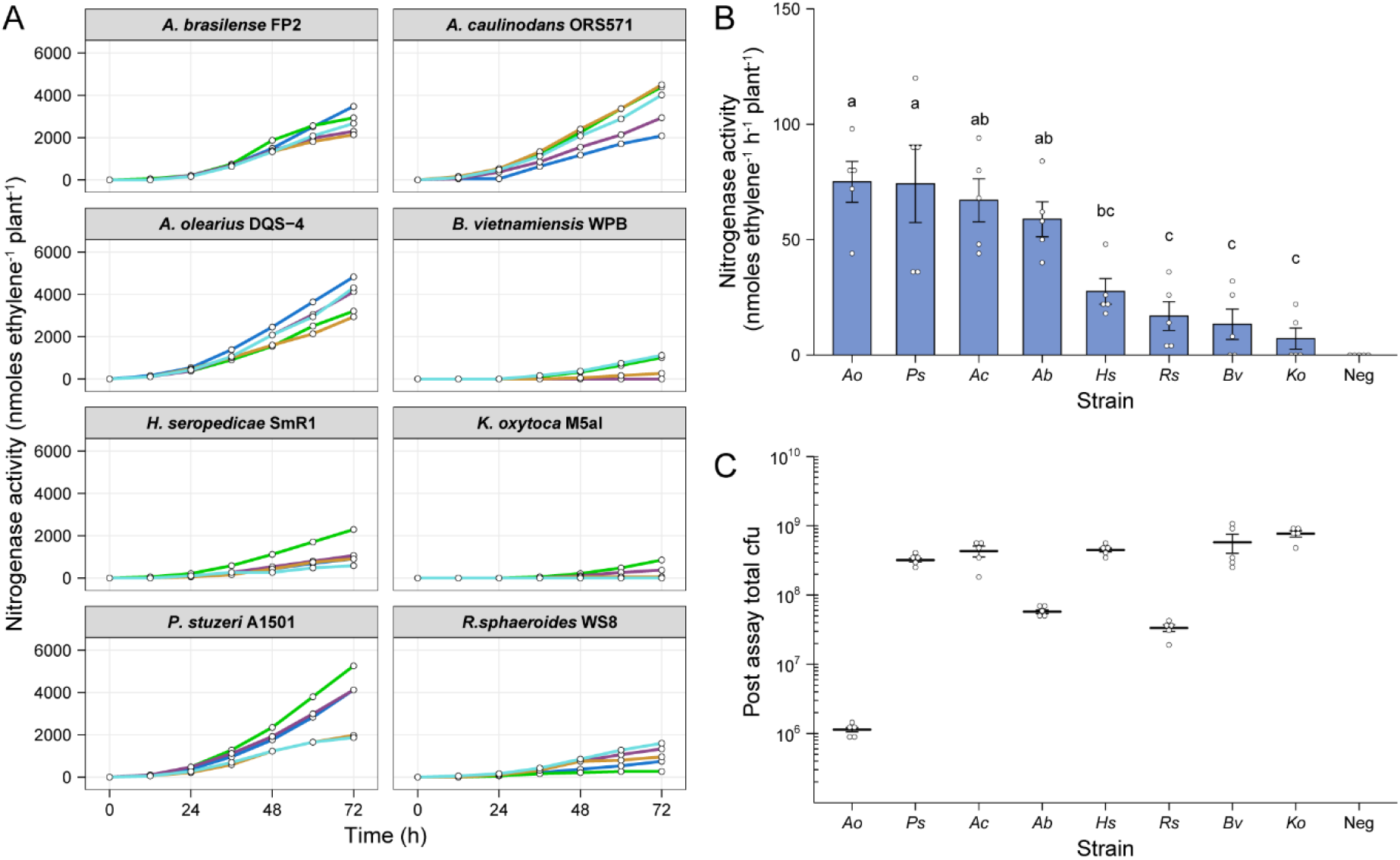
Comparison of associative nitrogen fixation between eight model bacteria. Assays were set up as described in Fig 2. Two millilitres of an OD_600nm_ 0.01 suspension of bacteria were used for inoculation and C_2_H_4_ production was measured at 12-h intervals over 72-h. **(A)** C_2_H_4_ production was measured for five biological replicates of eight genetically diverse diazotrophic bacteria inoculated gnotobiotically onto barley. **(B)** The mean ± s.e.m (error bars) rates of nitrogenase activity between 48-h and 62-h were calculated for the eight model diazotrophs. ANOVA and LSD tests with Bonferroni adjusted P-values were used to compare means. The negative uninoculated control omitted from these tests. Matching letters depict treatments that are not significantly different (P > 0.05) from each other but are significantly different (P < 0.05) from treatments with a distinct letter. **(C)** Mean ± s.e.m (error bars) of total cfu present in the assay systems as determined by viable counts after 72-h. Note that the low total cfu count for *Azoarcus olerarius* DQS-4 may be a result of poor viability of the strain upon re-isolation from they plant, as has been reported previously (17).

We also recovered bacteria from the assays and performed viable counts to estimate the total number of cfu. Interestingly, *Ao*, which exhibited the equal highest nitrogenase activity, was found to be the least abundant in the barley root systems (Fig 3C), indicating that individual cells may be capable of fixing a considerable amount of N relative to the other strains. However, we suspect that this result may be influenced by poor viability of the bacterium upon recovery from the plant, as has previously been documented (17). Conversely, the strains which were most abundant in the barley root system, *Bv* and *Ko*, exhibited the equal poorest nitrogenase activity, suggesting that these strains are well-adapted for proliferation in the barley rhizosphere, but poorly adapted to N fixation in this environment.

## Discussion

We have demonstrated that our simple *in planta* ARA is highly reproducible, rapid (can be completed in under two-weeks) and is applicable to a diverse range of diazotrophic bacteria. Thus, this assay represents a benchmark for quantification of associative bacterial N fixation.

The simple standardisation involved in our assay workflow (i.e. nitrogenase activity per plant) is one of its most beneficial features, offering a reduced workload compared to other potential approaches. We observed that the total population size of *Ac* in the assay system reached equilibrium after 9-dpi regardless of the starting inoculation density (Fig 2E), and that N fixation was consistent between these treatments (Fig 2C). Nevertheless, there was some variation observed in the total population size when comparing eight genetically diverse endophytes for N fixation (Fig 3). Therefore, in some instances it could be useful to standardise measurements of N fixation on per cell basis. We propose that fluorescent labelling of bacteria in combination with confocal microscopy or flow-cytometry (18) would be a suitable strategy if rates need to be expressed per cell (Fig 2E & Fig 3C) because some bacteria, such as *Ao*, exhibit poor viability upon recovery from plants (17) that could obscure measurements. Dual fluorescent reporter gene fusions could also be made to promoters of the nitrogenase structural gene *nifH* to assess the spatiotemporal dynamics of associative N fixation (19-22). This strategy has been useful in the past to detect *nifH* expression by *A*.*olerius* DQS-4 on the rhizoplane, endosphere and surrounding soil rice roots when carbon was supplemented into the soil (19, 20).

One of the major constraints of measuring associative N fixation is that optimal nitrogenase activity requires hypoxic conditions of ≤ 1% O_2_ (Fig 2C) which are detrimental for photosynthesis and plant growth (23). Although long-term exposure of plants to low oxygen ultimately results in anoxia, leading to acidosis and apoptosis, plants can postpone or even prevent tissue from becoming anoxic by tuning the expression or activity of energetically demanding metabolic pathways (24-26) and by producing non-symbiotic leghaemoglobins that help maintain redox status and remove reactive oxygen and nitrogen species (27). Remarkably, in our assay system, N fixation was stable over a 72-h period for all headspace O_2_ concentrations tested (Fig 2C), indicating that the plant is still able to provide adequate nutrients to fuel bacterial N fixation under these conditions. In addition to carbon, bacteria have been engineered to express N fixation genes in response to various plant-derived chemical signals, theoretically imparting upon them a degree of host-specificity for associative N-fixation (3, 8). Assessing such bacteria for associative N fixation using our assay system will be pivotal in the development of these strains, but it remains unclear as to whether plants can sustain production of chemical signalling molecules under low-oxygen stress.

In this work, we utilised barley as a host plant due to the highly uniform growth characteristics of seedings, but also due to its status as a model cereal for engineering the capacity for N fixation (https://www.ensa.ac.uk/) and the availability of a sequenced genome (28). We suspect that the assay could be readily extended to compare N fixation in other host plants, however this may require additional standardisation to account for differences in plant root mass. On the same note, the assay could be readily extended to assess the influence of various abiotic factors, such as plant growth substrates, nutrient levels, pollutants, temperature or light, or be used to explore the influences of abiotic factors on associative N fixation. This could be achieved for example by performing co-inoculation assays or assessing N fixation in non-sterile field soils, although this might be partially impeded by the presence of native N-fixing bacteria. Alternatively, defined synthetic communities of bacteria (29) could be inoculated as competitors for the diazotroph of interest. The validation and optimisation of our assay presented here has paved the way for such future extensions.

## Materials and Methods

### Preparation of inoculant strains for *in planta* ARA

Bacterial strains used in this study are listed in Table 1. *Klesbsiella oxytoca* was cultured on LB agar (30), *Burkholderia vietnamiensis* was cultured on TY agar (31) and the remaining strains were cultured on UMS agar (32) with 300 μM nicotinate, 10 mM NH_4_Cl_2_ as a sole nitrogen source, and either 30 mM malate (for *Azoarcus olearius, Azospirillum brasilense* and *Herbaspirillum seropedicae*) or 20 mM succinate (for *Azorhizobium caulinodans*) as a sole carbon source. All strains were grown at 28°C, except for *A. caulinodans* which was cultured at 37°C. For *in planta* ARAs, inoculants were prepared by streaking single colonies of bacteria onto agar slopes in 30 mL universal tubes. After 1-2 days incubation, cultures were washed from the slopes three times with PBS to remove residual N and resuspended in N/C-free UMS media at OD_600nm_ 0.001 – 1. The exact OD600_nm_ values are given in the results section for each experiment.

**Table 1.** Bacterial strains used in this study.

| Strain | Description | Origin |
| --- | --- | --- |
| *Azoarcus olearius* DQS-4 | Betaproteobacterium, isolated from oil-contaminated soil in Taiwan. | (33) |
| *Azorhizobium caulinodans* ORS571 | Alphaproteobacterium, isolated from *Sesbania rostrata* stems. | (34) |
| *Azospirillum brasilense* FP2 | Alphaproteobacterium, spontaneous St<sup>R</sup> mutant of Sp7 which was isolated from Tropical grasses in Brazil. | (35) |
| *Burkholderia vietnamiensis* WPB | Betaproteobacterium, isolated from *Populus* (cottonwood). | (36) |
| *Herbaspirillum seropedicae* SmR1 | Betaproteobacterium, spontaneous St<sup>R</sup> mutant of Z78 which was isolated from *Sorghum bicolor* in Brazil. | (37) |
| *Klebsiella oxytoca* M5a1 | Gammaproteobacterium, human pathogen isolated from soil. | (38) |
| *Pseudomonas stutzeri* A1501 | Gammaproteobacterium, isolated from the rice rhizosphere in southern China. | (39) |
| *Rhodobacter sphaeroides* WS8 | Alphaproteobacterium, isolated from soil in Ithica, NY. | (40) |

### *in planta* ARA protocol

*Azorhizobium caulinodans* ORS571 was used as a model strain for validation and optimisation of the *in planta* ARA. Golden promise barley seeds were initially surface sterilised by submersion in 70% ethanol for 2 min and 7% NaOCl for 2 min, then rinsed thoroughly in sterile water and germinated in the dark at room temperature for 2-days on 0.9% water agar. To house the barley seedlings, 130 mL Schott bottles were filled with 50 g of industrial grade sand 15 mL of N-free and C-free rooting solution (CaCl_2_•2H_2_0 2.67 mM, KCl 276 µM, MgSO_4_•7H_2_O 2.13 mM, Fe EDTA 26.67 µM, H_3_BO_3_ 93.33 µM, MnCl_2_•4H_2_O 24 µM, ZnCl_2_ 2.13 µM, Na_2_MoO_4_•2H_2_O 1.33 µM, CuSO_4_•5H_2_O 0.8 µM, KH_2_PO_4_ 1.33 g/L, Na_2_HPO_4_ 1.52 g/L), then autoclaved. One seedling was sewn into each bottle and immediately inoculated with 2 mL of bacterial suspension in UMS. The openings of Schott bottles were subsequently covered with sterile cling film and placed in a growth chamber with a 23 °C 16 h light / 21 °C 8 h dark cycle. At 6-dpi the Schott bottles were placed in a controlled atmosphere cabinet adjusted to 1% O_2_ by flushing with N_2_ gas, left for one hour and sealed with a rubber septum. To start the assay, ten percent of the headspace atmosphere (16.5 mL of air) was immediately replaced with C_2_H_2_ (13 mL) using a 20 mL syringe with needle and the plants were returned to the growth chamber. The evolution of C_2_H_4_ from C_2_H_2_ was measured at various timepoints outlined in the results section using GC-MS (Clarus 480 gas chromatograph, PerkinElmer) as previously described (41). Treatments for all experiments were performed with five biological replicates (i.e. five agar slopes of the inoculant and five Schott bottles containing barley seeds).

### Recovery of bacteria and estimation of population size

Bacteria were recovered from the root surface and soil of *in planta* ARA systems after 72-h in N-fixing conditions (9-dpi total) by adding 25 mL of PBS to the Schott bottles and vigorously agitating for 30 s. Viable counts were performed by establishing 10-fold serial dilutions of the resulting homogenous bacterial suspension from each Schott bottle and spotting 50 μL aliquots on non-selective agar plates. Colony morphology was examined to confirm that contamination had not occurred, and the total number of cfu present was estimated based on the total volume (60 mL).

### Statistical analysis

Total C_2_H_4_ production was calculated for each timepoint by deriving the fraction of the C_2_H_4_ peak area compared to C_2_H_2_, then multiplying this value by the number of C_2_H_2_ nmoles originally injected into the headspace based on the ideal gas law (5.31 x 10^5^ nmoles C_2_H_2_). Rates of nitrogenase activity were subsequently calculated between two time points as described in the Results section. All statistical analyses were performed using the agricolae and RStatix packages in R (42) and relevant information regarding each statistical test is provided in the figure captions.

## Acknowledgements

TH is the recipient of an 1851 Royal Commission for the Exhibition of 1851 Research Fellowship (RF-2019-100238) and Wolfson College, University of Oxford Junior Research Fellowship. This work was supported by the Biotechnology and Biological Sciences Research Council [grant numbers BB/L011484/1, BB/T006722/1].

